# GeneBench-Pro: Evaluating Multistage Statistical Reasoning in Genomics, Quantitative Biology, and Translational Biomedicine

**DOI:** 10.64898/2026.06.29.735386

**Authors:** Jeremy Li, Suyash Shringarpure, Edmund Wong, Andrew Ho

## Abstract

We introduce GeneBench-Pro, an expanded and improved version of GeneBench that comprises harder problems across a wider breadth of domains. GeneBench-Pro is a benchmark for AI agents performing realistic multi-stage scientific analyses in genomics, quantitative biology, and translational biomedicine which seeks to capture the complexity of real-world problems that computational life scientists face when tasked with producing a conclusion upon which a downstream scientific or translational decision is contingent. The benchmark comprises 129 evaluations targeting quantities of direct practical relevance across 10 primary domains and 21 terminal subdomains, with a genomics-centered core. Similarly to GeneBench, each problem provides the agent with brief context, a target estimand, and minimal guidance otherwise; the agent must then navigate multiple dependent decision points; *i.e.*, substantive inferential forks where a plausible wrong choice changes the downstream analysis, to identify and execute the correct analysis workflow and arrive at the correct answer. Relative to GeneBench, GeneBench-Pro adds 29 new problems, drops three, and introduces significantly redesigned versions of 54 of the remaining 100 overlapping problems. 82 of the 129 problems were reviewed by external domain experts, whose findings led to prompt/data modifications and redesign of those problems whose targets were not sufficiently identifiable. Ten externally reviewed problems are released publicly, 50 held-out problems were provided to Artificial Analysis for independent third-party model benchmarking, and the remainder are retained as an internal holdout. In evaluations over the full 129-problem suite, GPT-5.6 Sol reaches an eval-level pass rate of 28.7% at the `max` reasoning level, and GPT-5.6 Sol Pro reaches 31.5% in separately reported GPT Pro runs. GPT-5.5 reaches 12.0%, GPT-5.4 reaches 8.9%, and the strongest non-GPT baseline, Claude Opus 4.8, reaches 16.0%. As with GeneBench, models often complete substantial portions of the workflow but exhibit a consistent gap between *noticing* and *acting* by identifying local diagnostic signals but failing to propagate the implications to the corresponding analysis decision. As a result, models often select wrong estimators or persist on initially plausible but incorrect analysis paths. GeneBench-Pro therefore measures an emerging capability of long-horizon biological reasoning that remains unreliable.

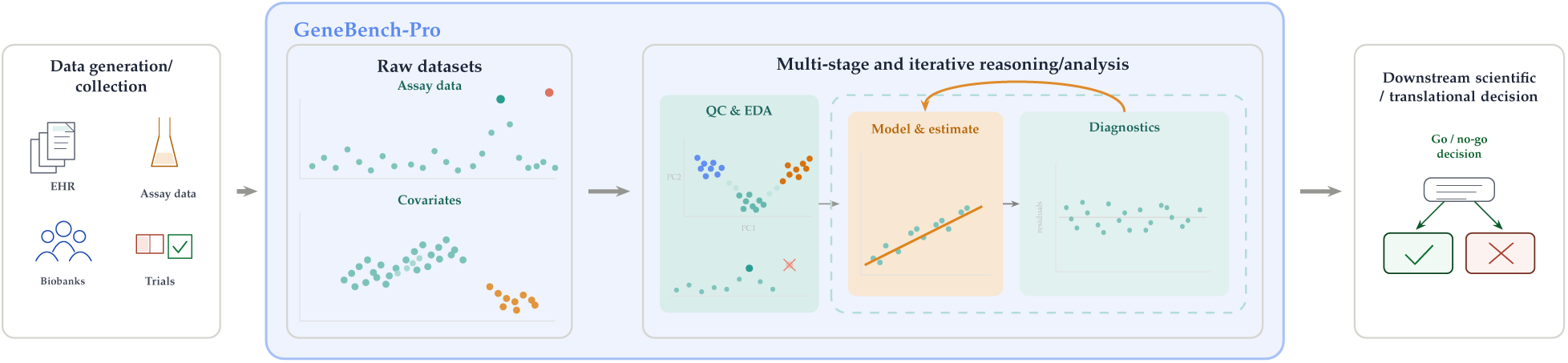

## Introduction

Agentic systems now perform strongly on basic software engineering evaluations, while newer suites such as SWE-Bench Pro, SWE-Lancer, DeepSWE, and FrontierSWE have shifted measurement toward longer-horizon, economically grounded, and deeper repository-level work; ^1–4^ broader evaluation efforts such as FrontierScience, FrontierMath, and Humanity’s Last Exam target difficult expert-level and novel-problem settings, ^5–7^ and METR’s recent time-horizon analyses likewise suggest that the duration of tasks frontier agents can complete autonomously is increasing rapidly. ^8^ Simultaneously, biology foundation models such as ESM3, Evo 2, and Omnii have pushed protein and genome modeling to new scale and fidelity. ^9–11^

Yet there has been relatively little formal examination of AI performance on the broader routine process that underpins much of modern life science research: executing a multi-step quantitative analysis starting from potentially errorful raw data, proceeding through a series of contingent procedures requiring statistical reasoning and strategic judgement, and ending at a decision-relevant conclusion (see **Figure 1B** for a high-level schematic of the typical process). This class of work is a major practical bottleneck in data-rich fields including genomics, proteomics, transcriptomics, and metabolomics; for example, recent reviews in the genomics literature argue that as sequencing has scaled, downstream computation and analysis, rather than data generation, have become the central bottleneck. ^12–14^

**Figure 1:**
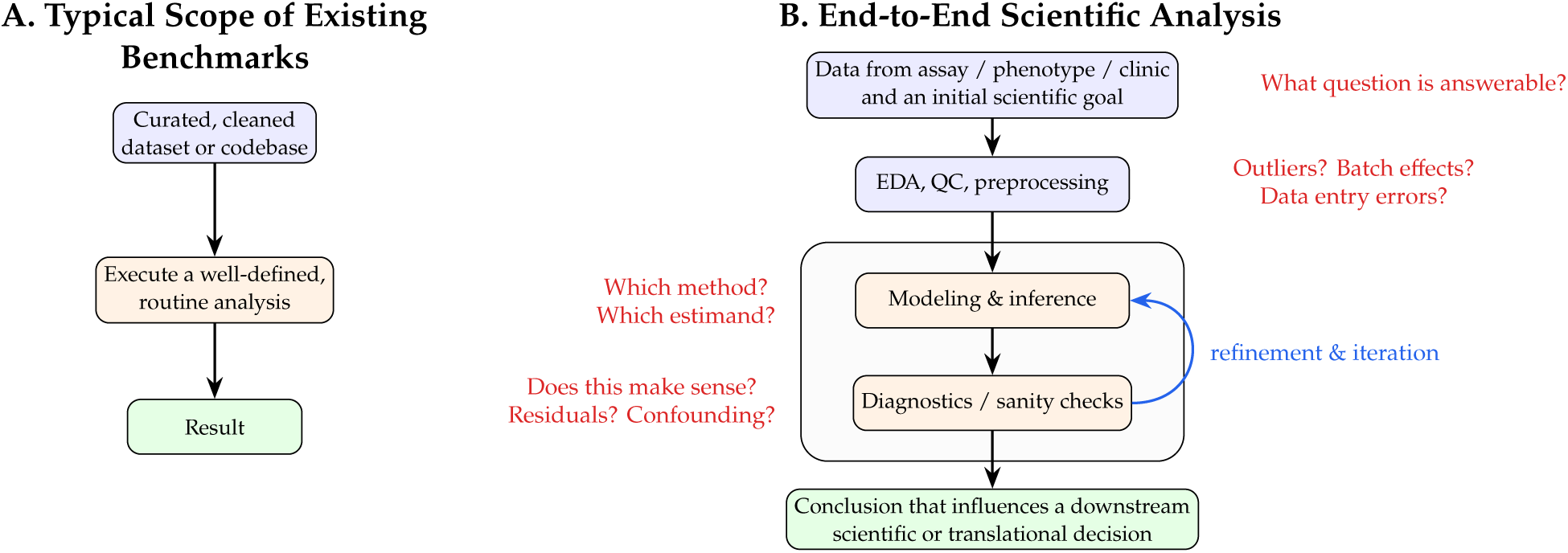
The benchmark gap. **(A)**: many existing biology AI benchmarks begin from a curated dataset and a highly specified prompt and evaluate a narrowly scoped analysis step with a cleanly verifiable answer. **(B)**: real-world scientific analysis more often spans a wider and more iterative process: data of unknown quality are obtained from some external source, and analysts must decide what questions are answerable, perform quality control and exploratory analysis, choose models and estimands, diagnose failures, and ultimately reach a conclusion that can influence the next scientific or translational decision. GeneBench-Pro is intended to evaluate this broader process rather than just its constituent components.

In contrast to most engineering tasks, scientific research is far more iterative, open-ended, and ambiguous. Its core challenges stem not from the execution of analytical workflows, but from the importance of scientific intuition or “research taste”: chains of judgment calls about what question the data can support, what data to include, which estimand or model is appropriate, whether diagnostics invalidate initial hypotheses, and when the evidence is strong enough to support a conclusion.

Older biology benchmarks mostly covered narrower forms of the workflow or emphasize breadth over depth (see **Figure 1A**), while newer efforts have begun moving closer to longer-horizon and agentic biology work. ^15–24^ Yet these still comprise fairly narrowly scoped problems that have been devised with respect to specific real-world datasets — problems which, even worse, often are not convincingly shown to only have one unique answer rather than a *distribution* of possible valid answers resulting from one of many possible defensible approaches, leading to difficulties in grading and uncertainty in whether ostensible failures should be taken at face value.

There therefore remains a gap in the literature for robust benchmarks which test whether agents can estimate quantities of biological interest from multi-stage analyses in which the accuracy of the final estimate is contingent upon accurate upstream statistical reasoning and diagnostic decision-making. GeneBench ^25^ was our first effort to address this gap directly, and leveraged simulated data, which enabled us to design sophisticated problems requiring multiple steps of contingent statistical decision-making while remaining confident in the fidelity of model grading.

Here we introduce GeneBench-Pro, an updated version of GeneBench comprising problems spanning industry and academically relevant domains that extends beyond the original scope of genomics into molecular and quantitative biology, pharmacogenomics, cancer biology, microbial genomics, clinical translation, and other settings where multistage statistical reasoning is required. As with GeneBench, each problem is a self-contained, multi-step analysis that provides (1) a realistic, messy dataset intended to reflect the data a scientist would receive from a lab, EHR system, or other collection pipeline, and (2) a *minimum viable prompt* which provides brief experimental context and defines a target estimand. That estimand is chosen to reflect a quantity that would inform a downstream decision in practice.

The current suite contains 129 evaluations across 10 primary domains and 21 terminal subdomains, with a genetics-centered core in population genetics, statistical genetics, quantitative genetics, and regulatory/molecular omics, and adjacent coverage in clinical genetics and pharmacogenomics, cancer somatic genomics and liquid biopsy, functional perturbation, proteomics, microbial genomics, and forensic genetics. GeneBench-Pro also adds a step of external scientific review, where the full design process, relevant files, and analytical setup for a given problem were exposed to an external domain expert to verify realism, scientific validity, and to solicit ideas on how to further harden the problem. This review process was performed for 82 of the 129 problems.

In evaluations over the full 129-problem suite, GPT-5.6 Sol reaches a 28.7% pass rate with `max` reasoning. At the highest performing reasoning level for each mainline GPT-family row, pass rate rises from 4.9% for GPT-5.2 to 8.9% for GPT-5.4, 12.0% for GPT-5.5, 16.5% for GPT-5.6 Luna, 23.3% for GPT-5.6 Terra, and 28.7% for GPT-5.6 Sol. Separately reported GPT Pro runs reach 8.5%, 16.3%, 20.5%, 23.6%, 28.5%, and 31.5% for GPT-5.2 Pro, GPT-5.4 Pro, GPT-5.5 Pro, GPT-5.6 Luna Pro, GPT-5.6 Terra Pro, and GPT-5.6 Sol Pro, respectively. Among the evaluated non-GPT models, pass rates range from 0.6% to 16.0%, with Claude Opus 4.8 the strongest non-GPT baseline. Manual examination of model-reported reasoning suggests that the main qualitative improvement in stronger models lies less in noticing the relevant diagnostic clues than in turning those observations into concrete corrective and model-selection decisions that move the analysis onto the correct path.

We first describe the scope of GeneBench-Pro using a high-level atlas of the problem space, introduce the main design constraints required to make this class of decision-heavy scientific analysis benchmarkable, and summarize the external-review process used to ensure that our problems are robust. We include an illustrative clinical genomics problem to make these design constraints concrete, then present benchmark-wide results and discuss qualitative improvements in model performance.

Of the 129 total GeneBench-Pro problems, we chose a subset of ten to open source. Another disjoint subset of 50 problems was chosen to serve as an external benchmark against which third parties may evaluate their own models via Artificial Analysis. The remainder remain an internal hold-out.

### Benchmark Scope and Construction

GeneBench-Pro, a collection of 129 problems across 10 primary domains and 21 subdomains, measures whether an agent can identify and execute the quantitative analysis required to estimate the target estimand from potentially errorful datasets with minimal guidance. **Figure 2** illustrates the domain coverage of the current suite.

**Figure 2:**
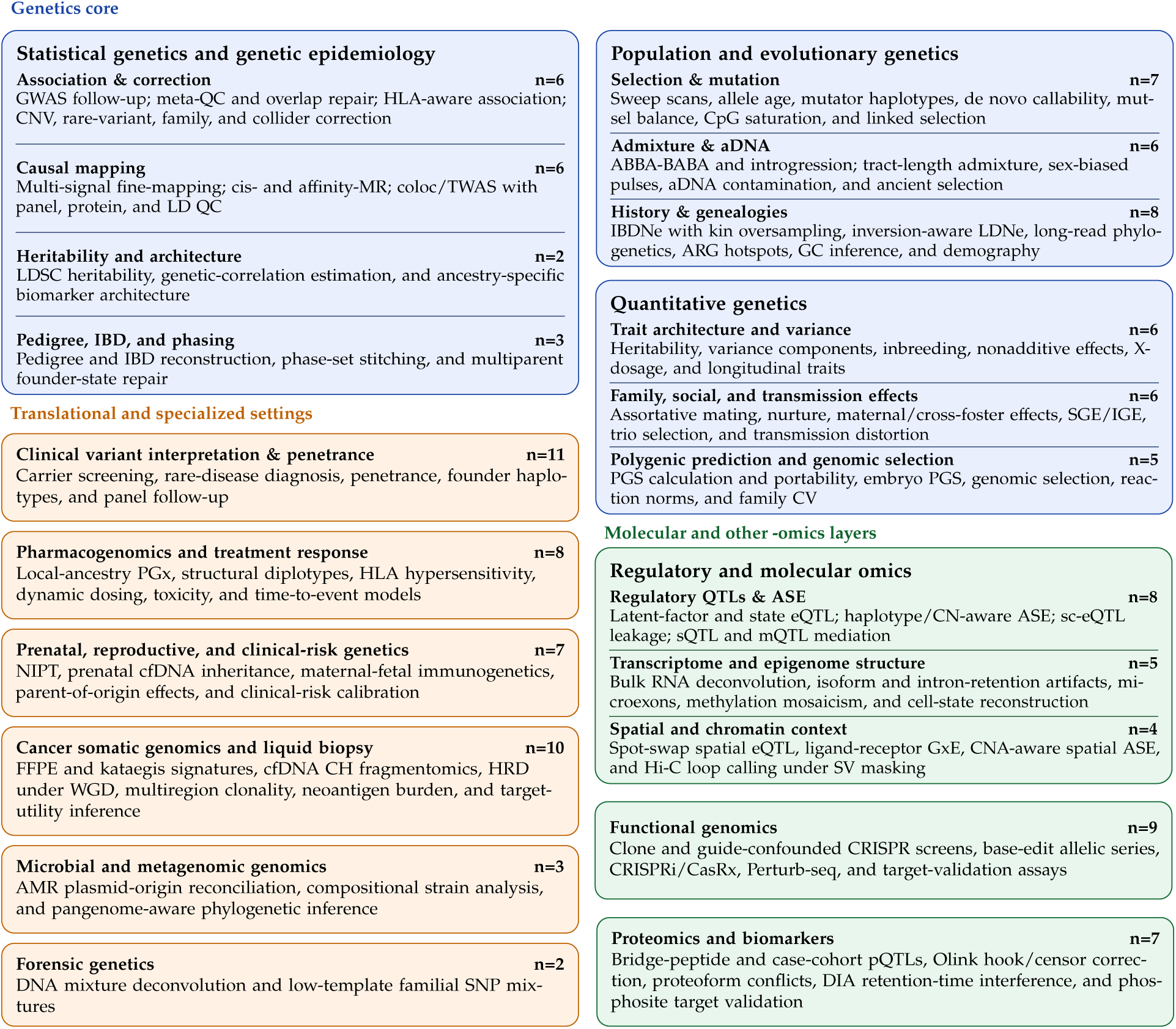
Domain atlas of the current GeneBench-Pro suite. GeneBench-Pro comprises 129 problems across 10 primary taxonomy domains and 21 terminal subdomains. Nested subcards expose the terminal subdomains within the larger domains. *Abbreviations:* GWAS, genome-wide association study; QC, quality control; HLA, human leukocyte antigen; CN, copy number; CNV, copy-number variant; SV, structural variant; MR, Mendelian randomization; coloc, colocalization; TWAS, transcriptome-wide association study; LD, linkage disequilibrium; LDSC, linkage disequilibrium score regression; IBD, identity by descent; PGx, pharmacogenomics; CpG, cytosine-phosphate-guanine; aDNA, ancient DNA; ABBA-BABA, four-taxon introgression test; IBDNe and LDNe, effective population-size inference from identity-by-descent and linkage disequilibrium, respectively; ARG, ancestral recombination graph; GC, gene conversion; PGS, polygenic score; SGE, social genetic effects; IGE, indirect genetic effects; QTL, quantitative trait locus; ASE, allele-specific expression; eQTL, expression quantitative trait locus; pQTL, protein quantitative trait locus; mQTL, methylation quantitative trait locus; sQTL, splicing quantitative trait locus; sc-eQTL, single-cell expression quantitative trait locus; CRISPR, clustered regularly interspaced short palindromic repeats; CRISPRi, CRISPR interference; CasRx, an RNA-targeting CRISPR effector; Hi-C, genome-wide chromosome conformation capture; GxE, gene-by-environment interaction; CNA, copy-number alteration; NIPT, noninvasive prenatal testing; cfDNA, cell-free DNA; FFPE, formalin-fixed, paraffin-embedded; CH, clonal hematopoiesis; HRD, homologous recombination deficiency; WGD, whole-genome doubling; AMR, antimicrobial resistance; SNP, single-nucleotide polymorphism; DIA, data-independent acquisition.

Across the benchmark, an agent must filter and correct data, identify QC or ascertainment problems, choose methods, perform statistical inference, revise the analysis when intermediate results disagree with the initial plan/hypothesis, and produce a final quantitative answer. Many problems are framed as decision points in genetics-backed drug discovery and translational research, such as whether a GWAS signal survives correction strongly enough to advance and which gene or protein should be nominated as the likely effector target, while others are framed around more academically oriented questions, such as whether an observed pattern is better explained by selection or demography and which pedigree, haplotype, or ancestry reconstruction is supported by the data.

GeneBench-Pro preserves the same problem setup as GeneBench—a minimum viable prompt, agent-visible staged data files, and a defined output schema, but enhances the difficulty of the problems, expands the problem set, and shifts more of the difficulty away from addressing messy data toward decision-heavy statistical reasoning. Specifically, we began with GeneBench’s 103 problems—we then withdrew three due to fatal issues identified; of the remaining original hundred, we significantly redesigned and hardened 54; we then added 29 new problems, resulting in a total of 129 problems.

Relative to GeneBench, GeneBench-Pro also increases coverage in clinical and translational genomics, pharmacogenomics, liquid biopsy, proteomics, microbial genomics, and specialized molecular-omics settings, and includes extensive external scientific review of 82 problems in the full suite (described below).

### Public Case Studies

For each of the ten public GeneBench-Pro packages, we provide, in addition to the data and prompt for each problem, detailed reports including a description of the data generating process, an analysis walkthrough corresponding with the correct answer, validation and ablation evidence, and general construction rationale. These are distributed with the Hugging Face dataset.

### Benchmark Setup

Each GeneBench-Pro problem is packaged as a self-contained scientific analysis. The agent receives an isolated workspace containing a *minimum viable prompt*, staged files, and a standard scientific Python stack. The prompt specifies the scientific question/task and target estimand without explicitly prescribing the workflow to be executed. The files are intended to resemble what an analyst might actually receive from assays or clinical systems rather than cleaned toy datasets. Each problem involves a chain of dependent decision points such that an incorrect choice at any stage propagates into downstream errors and ultimately failure to recover the final correct target.

The agent operates in a realistic sandbox with the staged files, access to general-purpose scientific libraries including `numpy`, `pandas`, `scipy`, `scikit-learn`, `statsmodels`, `lifelines`, `matplotlib`, and `seaborn`, and standard genomics bioinformatics tooling such as PLINK 2.0, `pysnptools`, `bed-reader`, `bedtools`, `pysam`, etc. (see **Methods**)—though the problems are designed not to require or rely on access to these tools. **Supplementary Figure 1** shows a schematic of the agent environment. Success therefore depends both on the agent recovering the analysis from the data as well as accurate implementation of the relevant methods.

### Construction, Validation, and Grading

Open-ended scientific analysis is difficult to benchmark precisely because real data often admit multiple defensible analysis choices. For example, QC thresholds, model parameterizations, and reporting conventions can vary across analysts without there being only a single unambiguously correct analytical choice. If the outcomes of a benchmark change because one agent uses one defensible cutoff or convention while another agent uses a different, yet equally defensible one, this might reflect the arbitrary nature of that benchmark’s design choices rather than the quality of scientific reasoning.

This is a particularly underappreciated issue in many existing long-horizon biology benchmarks, which are often based on real historical datasets around which new, multi-step questions are devised. In such data, extreme ambiguity is often present, to the extent that if one is, for example, tasked to apply a three-step QC or modeling pipeline to some dataset, there is rarely necessarily exactly one correct path through this garden of forking paths — at each of these three steps, there is a substantial probability that there exists an equally defensible choice that was not considered by the benchmark designer that ultimately leads to agents which make this choice to be graded as a failure, leading to a naturally decaying terminal pass rate with the length of the required inferential chain just by virtue of real-world messiness.

This can then lead to silent failure modes in which a benchmark may appear difficult due to the contribution of this mode of “failure” to overall benchmark performance, making it difficult to evaluate exactly what it is that such benchmarks are measuring. A useful benchmark for this type of work must therefore be insensitive to nearby defensible analyst choices, but sensitive to missing scientifically necessary stages.

In order to implement these multi-stage (“cascaded”) setups in a way where we can be confident that a “fail” actually represents a scientific failure and not a result of the aforementioned natural attrition, GeneBench-Pro problems are based on constructively simulated problems where the full causal structure is known and where we simulate the full data-generating process (DGP). Simulation allows us to directly tune the complexity and difficulty of this cascade while ensuring that (1) QC-sensitive decisions are robust to small researcher-choice variation, (2) plausible wrong analyses fail for substantive reasons, and (3) the graded endpoint is actually recoverable from the agent-visible data.

We quantify this cascaded structure through the number of *decision points* in each problem: substantive inferential forks where a plausible wrong choice leads to a qualitatively different downstream answer. The number of these decision points ranges from 3 to 13 across the current suite (with a median of 6).

Operationally, problem development begins from a real-world analysis pattern and a target estimand. These real-world analysis patterns are synthesized from the literature and domain expertise to reflect common, high-impact scientific questions and workflows, and are specifically chosen so they do not recapitulate well-known textbook examples or papers, so as to avoid the risk of benchmarking against memorized solutions. Data are then simulated so that the correct answer is recoverable from the staged files (for example, the maximum likelihood estimate of a parameter resulting from the correct approach would be considered as the ground truth value for grading, rather than the parameter under which the data were generated). A minimum viable prompt containing the minimum amount of information required to make the correct answer identifiable is then constructed. Prompts also include a standardized instruction that the data came from a real experiment, analytical reasoning quality is graded alongside numerical correctness, and the final response must be exactly one JSON object.

Once an initial draft of a problem is completed, extensive validation is performed. Results from analyses involving plausible but incorrect decisions at the various inference stages are checked via ablation and verified to be sufficiently distinct from the graded answer. Independent reviews for scientific validity, methodological soundness, and target identifiability are conducted in order to ensure that the evaluation is testing the intended capabilities rather than whether agents can infer an artifact-specific workflow preference that is not uniquely supported by the data. Problem drafts are then iteratively audited through multiple rounds of frontier-model pilots and detailed trace analyses in order to check for unintended leakage, alternative unintended pathways to the correct answer, prompt-grader mismatch, and robustness. This process is intended to ensure that wrong-but-plausible analyses fail for substantive reasons and that passing runs reflect the intended inferential path rather than shortcuts. **Table 1** summarizes the main benchmark-level constraints that follow from these requirements.

**Table 1:** Primary design constraints in GeneBench-Pro. Together, these are intended to keep the graded endpoint scientifically identifiable while preserving realistic ambiguity and data messiness.

| Principle | Benchmark requirement | Failure mode if violated |
| --- | --- | --- |
| *Ground truth and identifiability* |  |  |
| Recoverable target | Agents are graded on recovering the quantity that is actually recoverable from agent-visible data, and not the hidden data-generating parameters. | Correct analyses can be marked wrong because the parameter under which data were generated is unrecoverable (e.g., due to sampling variation in the DGP). |
| Unique, identifiable answer | The staged evidence along with a minimum viable prompt supports one uniquely defensible answer. If multiple approaches would naively appear reasonable, the data or prompt contain some empirical constraint that reasonably rules out all but one. | The task becomes under-specified, and success depends more on workflow preference rather than reasoning from the available evidence. |
| Clear numerical separation from incorrect answers | A comprehensive ablation suite demonstrates that plausible but incorrect analyses and shortcut methods yield materially different answers and fail by clear margins. | Wrong analyses can be graded as correct. |
| *Problem specification* |  |  |
| Minimal viable prompt | The scientific question and graded estimand(s) are clearly defined, but the prompt does not give prescriptive instructions on how to perform the analysis. | The task either turns into evaluating how well agents can execute a well-defined analysis, or leaves multiple defensible interpretations of how the final answer should be estimated. |
| Robust QC requirements | When QC is required as part of reaching the solution, nearby reasonable thresholds lead to the same graded outcome. | The benchmark measures arbitrary cutoff choice rather than recognition of the qualitative problem. |
| *Scientific workflow fidelity* |  |  |
| Fully simulated problems | Simulating data allows us to tune each aspect of the data-generating process such that realism, multi-stage inference, effect sizes, and diagnostic clues can be precisely controlled. | Difficulty, answer separation, and correctness may become difficult to calibrate. |
| Multi-stage inference | Problems require navigating ordered inferential forks in which upstream filtering, representation, and statistical model choices materially affect the final graded endpoint. | The benchmark reflects smaller, self-contained units of end-to-end analysis rather than the full scientific workflow. |
| Representation and nuisance variable discovery | Problems require recovery of the relevant scale, orientation, harmonization state, selection or censoring process, batch or latent-group structure, and QC-valid sample set before final estimation. | A statistically valid final model is applied to the wrong data or population, on the wrong scale, or on the wrong conceptual level. |
| Ablation-supported workflow adjudication | Plausible expert workflows are evaluated through ablations and sensitivity analyses, separating reasonable alternatives from choices that alter an identifiable stage of the inferential chain. | Success depends on an unreported analyst preference rather than a recoverable scientific target. |
| Observed failures occur across multiple decision points | Trace analyses display agent failures across multiple decision points rather than clustering around only a single decision point. | The problem effectively evaluates how well agents navigate a single difficult step rather than multiple difficult steps. |
| Literature-defensible solution | The intended correct solution involves standard or otherwise well-supported methods. | Success depends on benchmark-specific ad-hoc methods or unsupported analysis choices rather than well-established methods. |

GeneBench-Pro uses binary grading against recoverable targets under calibrated tolerances chosen to allow for numerical and implementation-level variation; the evaluation setup and package-level grading protocol is summarized in the **Methods**.

### External Scientific Review

External review was used to assess a subset of candidate problems in order to gain broader confirmation of the scientific defensibility of the problems as well as their adherence to the stated benchmark design principles.

Reviews focused on target identifiability given the agent-visible prompt and data, method implementations, realism, and estimator choices. Feedback was adjudicated against the benchmark design constraints outlined in **Table** 1. This review layer also reinforced the benchmark’s focus on statistical reasoning within quantitative biology: defining identifiable estimands, choosing among defensible finite-sample targets, checking method equivalence, respecting assay-specific artifacts, and recognizing when a realistic data-generating assumption changes the scientific answer.

#### Review Process

To limit exposure of unreleased tasks, reviewers received access to private repositories with their assigned problems. Each repository contained the prompt and data for each problem, a detailed problem report fully describing the problem design, validation materials, and an issue thread for their feedback.

The reviewer pool included 11 domain experts spanning graduate students, postdoctoral researchers, industry scientists, and professors. In total, 84 candidate problems received external review. 82 are included in the 129-problem evaluation, and two reviewed candidates were withdrawn due to fatal issues. The ten publicly released GeneBench-Pro problems were selected conditional on external review.

#### Review Themes and Resulting Changes

Table 2 gives representative excerpts from the reviews. For those problems where issues were identified via review, critiques tended to fall in a few categories: (1) the presence of hidden assumptions or ambiguity in estimand definitions which would result in non-identifiability of the graded target, (2) errors in the method implementations corresponding with the “correct” approach which would lead to correct answers being graded against an incorrect ground truth, (3) contrived setups which were nonrepresentative of real-world workflows.

**Table 2:** Examples of review findings. Short quoted excerpts from external-review comments followed by brief context explaining the scientific concern are shown with the resulting changes that were made.

| Finding type | Problem | Reviewer concern and context | Disposition or change |
| --- | --- | --- | --- |
| Underspecified estimand | Cis-MR with winner’s curse and LD | “the exact graded solution rests on a particular pipeline not uniquely dictated by the prompt.” The concern was that broad robust-MR, colocalization, or external-annotation strategies could be scientifically reasonable even if they did not match the specified ground truth pipeline. | Specified the residual-pruned finite-sample two-coefficient estimand in the prompt and answer contract. |
| Alternate defensible method failing | Recent-pulse sex-biased admixture | “I couldn’t understand what the posterior represented” and whether a solver should “propagate uncertainty into the final solution.” The reviewer identified a defensible posterior-weighted ancestry-fraction variant rather than a pure hard-call tract summary. | Allowed posterior-weighted ancestry fractions in grading, rather than requiring a hard-call result. |
| Method parity violation | Genetic correlation estimation via LDSC | The reviewer wrote that the ground truth was “not implementing actual LDSC” and that the critical issue was “wrong M, OLS not WLS.” The concern was that the benchmark was grading an ad hoc LD-score estimator while presenting itself as an LDSC genetic-correlation task. | Fixed the reference implementation to implement LDSC properly. |
| Underspecified prompt | Somatic TXR1 target activation | “SV-mediated but that’s not clear in the agent provided materials.” Private materials made the causal SV mechanism clear, but the solver-facing materials left room for interpreting high copy number without an SV as activation. | Added “SV-driven” wording to the solver-facing target definition. |
| Ground truth not recoverable | Dynamic PGx treatment response | The reviewer recommended changing the “ground truth to the DGP’s realized data” and noted that the validator was “suboptimal, a little ambiguous.” Several calibration, toxicity, and biomarker nuisance specifications were defensible under the original prompt. | Aligned the ground truth with the realized simulated data and made the validator target and nuisance specifications explicit. |
| Non-identifiable phasing | Copy-number-aware phased ASE | The reviewer argued it was “not possible to have a fixed polarity” for arbitrary haplotype labels and that the flanking-marker step could not be used “in its current state.” Marker relevance, copy-number context, and tumor purity were under-specified. | Redesigned the prompt and data with donor-specific phase blocks, explicit copy-number segments, marker relevance, and tumor-purity context. |
| Conflicting assay evidence | Long-read pseudogene phase diagnosis | “short reads and long reads” gave contradictory phasing evidence. The reviewer argued the target should state that phase posterior estimation should be based on long-read evidence, with QC used as pass/fail rather than as a continuous probability weight. | Specified long-read-only phase evidence and made phase-set quality a pass/fail QC screen. |
| DGP method implementation error | Assortative mating and indirect genetic effects | The reviewer found that the simulated child-parent PGI correlation “came out to be 0.38. This is too low” and traced it to a “faulty formula.” The review also noted that the solver-visible heritability parameter should be $h_f^2$ rather than the originally exposed quantity. | Rebuilt the simulator around the corrected child/mother/father PGI covariance, exposed $h_f^2$, and updated the validator equations. |

Corrections were proportional to the finding: minor issues led to prompt clarifications or surgical changes to the data files, whereas more substantive issues led to more substantial work in modifying the DGP, ground-truth generation, and in extreme cases, complete redesign or withdrawal of the problem. After corrections were applied, the standard loop of agent pilots *→* trace examination *→* further revisions was repeated to generate the final candidate.

### Illustrative Problem: DRX1 Carrier-Screening Residual Risk

Figure 3 illustrates a GeneBench-Pro problem modeled on carrier screening for a recessive genetic condition. DRX1 is the disease gene in this synthetic screening scenario: an individual is at reproductive risk only if both biological parents carry a reportable DRX1 allele. The agent is given raw carrier screening data and must estimate how much carrier risk remains after a negative screen, then combine that residual risk with the carrier frequency of potential reproductive partners.

**Figure 3:**
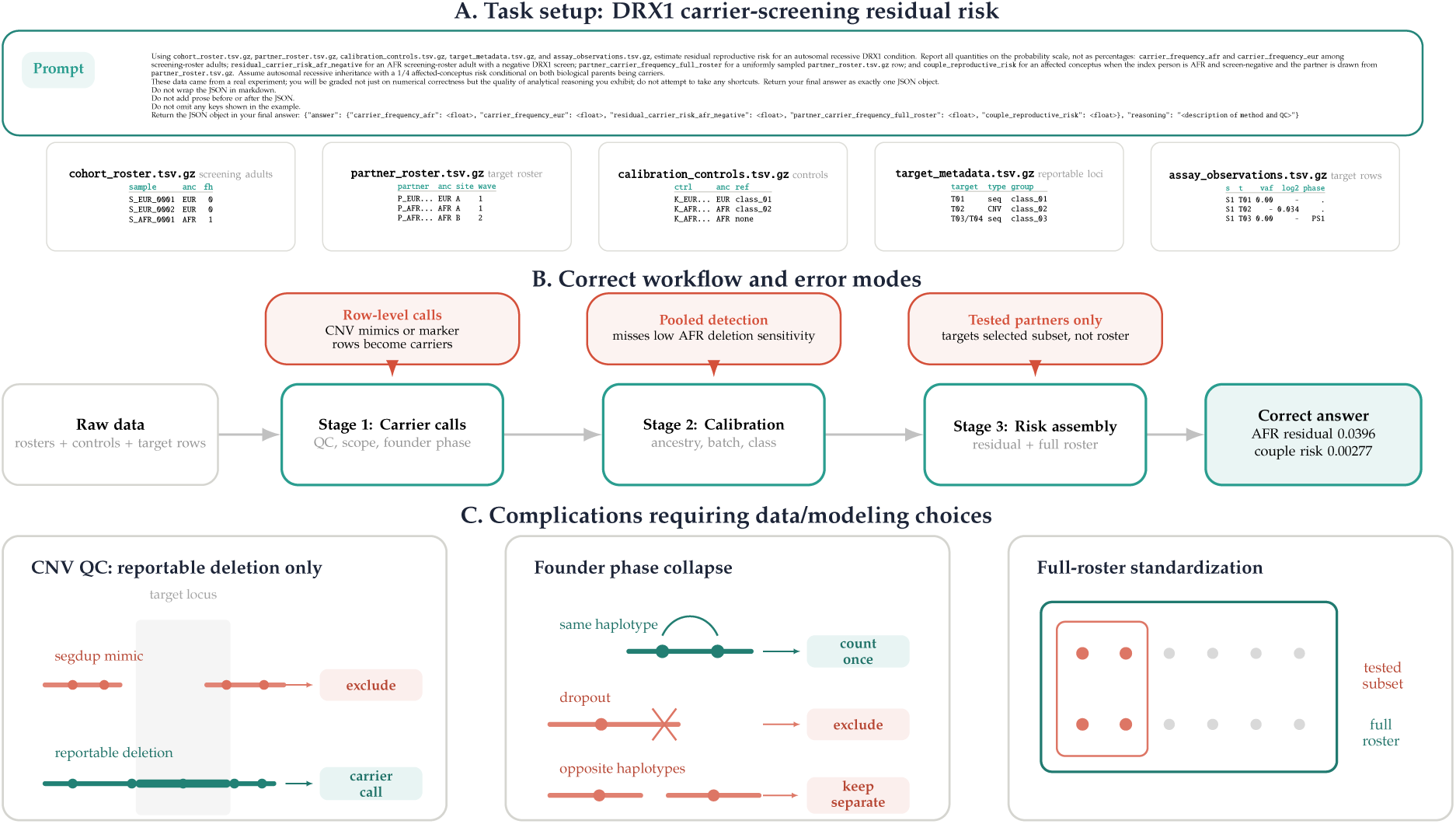
Illustrative GeneBench-Pro problem from clinical genomics: residual risk in carrier screening. The DRX1 problem asks an agent to estimate residual reproductive risk after a negative carrier screen using five staged tables: screened adults, potential partners, calibration controls, target metadata, and assay observations. **(A)** Problem setup: the agent receives a minimum viable prompt defining the probability-scale carrier, residual-risk, partner-frequency, and couple-risk targets; previews of the data files are also shown. **(B)** The answer requires navigating three linked stages: translating assay observations into reportable carrier calls, estimating assay performance from calibration controls, and combining residual carrier risk with the carrier frequency of the full partner roster. Red nodes mark shortcut analyses that resolve only part of this chain and which ultimately lead to the wrong final answer. **(C)** Cartoon of a few of the answer-changing decision points: separating reportable copy-number calls from segmental duplicate mimics, collapsing phased founder markers when they represent one haplotype, and standardizing partner risk to the full roster rather than only the tested subset.

The dataset has five tables. One table lists screened adults, including ancestry and family-history information. A second lists the full roster of potential partners: some partners have completed assay data, but the estimand is the average carrier probability over the entire roster, not just over the tested subset. A third table contains calibration controls that reveal how often each assay signal detects true carrier states or produces false positives. The final two tables describe the DRX1 assay targets and give the raw per-sample assay measurements.

The difficulty here is that the answer cannot be naively estimated as the raw carrier rate. The correct analysis must instead first correctly identify reportable DRX1 carrier classes, including an exon-level deletion while distinguishing copy-number mimics, properly deal with phased founder markers and whether to consider them a single haplotype, estimate assay sensitivity and false-positive rates by several covariants, compute residual carrier risk after an AFR negative screen from class-specific missed-carrier probabilities, and standardize partner carrier frequency to the full partner roster rather than to the selected tested subset. This example therefore illustrates a central GeneBench-Pro design goal: a minimum viable prompt and clearly defined target estimand are provided to the agent, but success ultimately depends on the agent recovering a multistage quantitative analysis path from available data rather than simply implementing a prescribed workflow.

## Results

We evaluated GeneBench-Pro on the full 129-problem suite across 60 evaluated model configurations spanning GPT-5.2, GPT-5.4, GPT-5.5, GPT-5.6 Luna/Terra/Sol, corresponding GPT Pro variants, and non-GPT baselines from Claude, Gemini, Grok, GLM, Kimi, DeepSeek, MiMo, Tencent, MiniMax, and Qwen. **Figure 4** summarizes the resulting unweighted per-problem pass rates, and **Supplementary Table 2** reports release-subset and review-status strata.

**Figure 4:**
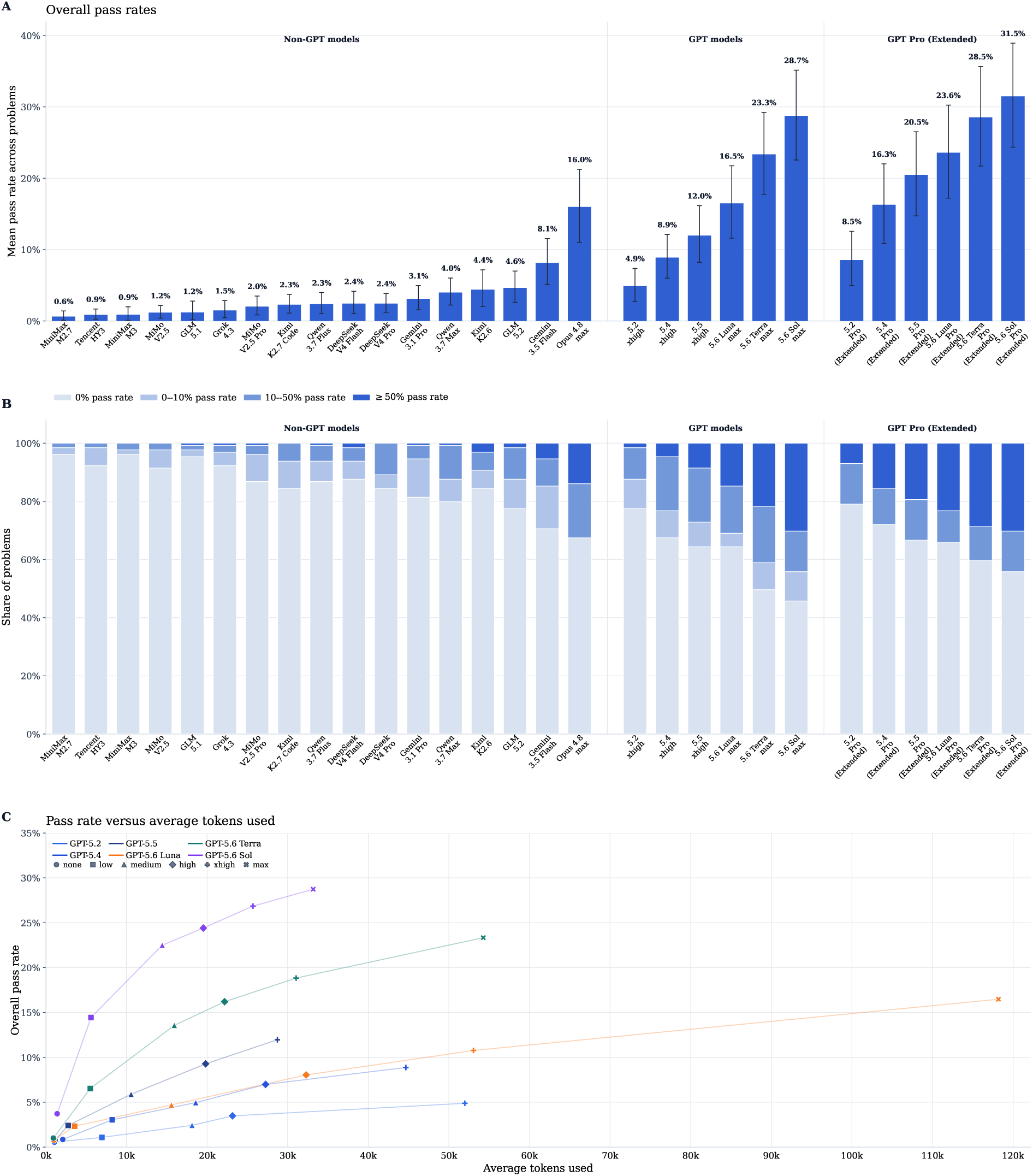
Performance across evaluated models. **(A)** Overall pass rate for headline rows, defined as the unweighted mean of per-problem pass rates across the 129 GeneBench-Pro problems. For models with multiple reasoning settings, panels A and B show the highest-pass-rate setting; the complete 60-row result table is reported in **Supplementary Table 1**. Error bars show 95% hierarchical bootstrap confidence intervals from 20,000 resamples, resampling problems and repeated runs within each problem. **(B)** Distribution of per-problem pass rates across four regimes for each headline row: 0%, 0–10%, 10–50%, and at least 50%. **(C)** GPT-family pass rate as a function of average tokens used, computed from model trace and response tokens under the internal GPT evaluation harness. GPT Pro (Extended) and non-GPT rows are shown in panels A and B and omitted from panel C because comparable token accounting was unavailable.

### Overall performance and unsolved tail

Overall pass rates remain low, indicating that the expanded and hardened suite is still far from saturated. At the best-performing reported reasoning level for each mainline GPT-family row, the unweighted mean pass rate rises from 4.9% for GPT-5.2 (`xhigh`) to 8.9% for GPT-5.4 (`xhigh`), 12.0% for GPT-5.5 (`xhigh`), 16.5% for GPT-5.6 Luna (`max`), 23.3% for GPT-5.6 Terra (`max`), and 28.7% for GPT-5.6 Sol (`max`). The separately reported GPT Pro runs reach 8.5% for GPT-5.2 Pro, 16.3% for GPT-5.4 Pro, 20.5% for GPT-5.5 Pro, 23.6% for GPT-5.6 Luna Pro, 28.5% for GPT-5.6 Terra Pro, and 31.5% for GPT-5.6 Sol Pro. The evaluated non-GPT rows range from approximately 0.6% to 16.0%, with Claude Opus 4.8 the strongest non-GPT baseline. Within the GPT-family models, increasing the reasoning level has a large effect: GPT-5.6 Sol rises from 3.7% at `none` to 14.4% at `low`, 22.5% at `medium`, 24.4% at `high`, 26.8% at `xhigh`, and 28.7% at `max` (**Figure 4C**).

A substantial unsolved tail remains (**Figure 4B**). Along the best mainline GPT-family rows, the share of problems with 0% pass rate declines from 77.5% for GPT-5.2 to 67.4% for GPT-5.4, 64.3% for GPT-5.5, and 45.7% for GPT-5.6 Sol, whereas the share reaching at least 50% rises from 1.6% to 4.7%, 8.5%, and 30.2%. The benchmark therefore remains dominated by hard items even in the strongest mainline row, but stronger models move a larger fraction of problems from the all-fail floor into partial or frequent success. Exact values underlying **Figure 4** and the corresponding numbers of valid samples per problem are reported in **Supplementary Table 1**.

Because release visibility and external-review status may introduce selection effects, we report descriptive pass rates for the locked result-reporting subsets and review strata (**Supplementary Table 2**). These strata were not randomized. In this evaluation, the Artificial Analysis subset had lower pass rates than the full suite for most reported rows, including the strongest GPT-family rows.

### Inferential chain length and action on intermediate diagnostic evidence

The number of decision points remains a useful concept in GeneBench-Pro: each problem is designed so that several upstream choices must be made correctly before the final estimand is estimable. In the current evaluation, we use these decision-point counts as qualitative metadata rather than as a quantitative stratification of model performance.

Manual review of model-reported reasoning from selected GPT-5.5/GPT-5.6 Sol comparisons suggests a consistent mechanism behind this scaling. In many failures, the agent notices the relevant local diagnostic clue but treats it as a local data cleaning issue rather than as evidence that should change the downstream statistical method and QC pipeline. **Table 3** shows representative excerpts from these comparisons. In these examples, the weaker model typically realizes that an initial correction is required, but then fails to sufficiently change its downstream analysis plan to achieve the final correct answer. In contrast, the stronger model is more likely to carry the same diagnosis through the inferential chain and consider its potential impact on subsequent methodological choices.

**Table 3:** Representative excerpts from the model-reported reasoning underlying responses from selected GPT-5.5 and GPT-5.6 Sol runs. In each case, both models identify or note the relevant local signal, but GPT-5.6 Sol more often uses that signal to correctly adjust its downstream methodology used to generate the final answer.

| Problem | GPT-5.5 pattern | GPT-5.6 Sol pattern |
| --- | --- | --- |
| **Pharmacogenomic time-to-event response with time-varying treatment:** treatment initiation, genotype-specific response, delayed pharmacodynamics, prevalent-user flags, and longitudinal biomarkers jointly determine the causal survival estimand. | **Handles treatment timing with a conventional Cox outcome model but does not address treatment-confounder feedback.**<br>"Fit a counting-process Cox model with treatment as a time-varying exposure, effective only after treat_start+90 days ... The model included G, treatment×G, baseline severity, age, and sex." | **Uses a more appropriate causal inference method to account for treatment-confounder feedback**<br>"Used a new-user marginal structural Cox model: excluded 818 flagged prevalent users, modeled treatment initiation with stabilized inverse-probability weights using baseline covariates and current biomarker, and treated exposure as time-varying with a 90-day efficacy lag." |
| **Conditional cell-type heritability with LD-score reference choice:** the analysis requires an effective sample-size calculation, reference-panel matching, summary-statistic QC, outlier-region removal, and a joint annotation model before enrichment can be interpreted. | **Applies basic allele-frequency and LD-region filters but selects the reference panel by concordance only.**<br>"After merging files, I kept GWAS maf≥0.05 and removed the BED LD-outlier regions, leaving M=13100. Panel A was selected by post-QC GWAS/ref allele-frequency concordance." | **More carefully defines the reference panel using comprehensive frequency, imputation-quality, association-statistic, and LD-region filters**<br>"SNPs were merged by ID and position, restricted to GWAS MAF≥0.05 and INFO≥0.9, filtered at $\chi^2 \leq$ 80, and excluded if inside the supplied BED regions, leaving M=12658." |
| **Bridge-calibrated peptide pQTL with genotype-dependent assay artifacts:** bridge-injection QC, plate-by-peptide normalization, peptide-specific genotype artifacts, and peptide roll-up jointly determine the additive protein-abundance effect. | **Adjusts for statistical artifacts but averages all six peptides, including those with discordant dose responses.**<br>"Verified 192 cohort samples with complete six-peptide measurements. Excluded four bridge injections with high erythro_score ($>0.1$). From the remaining bridges, estimated centered plate/peptide offsets, corrected sample log2 peptide intensities, averaged the six corrected peptides per sample, then fit OLS: log2 protein abundance ∼ dose + age + sex." | **Correctly uses dose-response consistency to exclude discordant peptides.**<br>"Verified 192 unique samples with complete six-peptide data and matching plates. Excluded four bridge runs with extreme erythro_score ($>0.1$), then normalized each peptide by its plate-specific clean-bridge mean. Peptide-specific dose slopes flagged pep_02 and pep_04 as discordant; the remaining four peptides were averaged." |

## Discussion

Agentic abilities in software engineering, computer use, broad scientific reasoning, and general capabilities have been increasing at a rapid pace, as evidenced by recent model progress and benchmark turnover across long-horizon software engineering and adjacent skills. ^1–4,7,26^ In parallel, genomic and biomedical benchmarks have expanded from knowledge and computational-biology evaluations toward more realistic biological workflows. ^20,27^ Recent biological benchmarks have begun to probe longer-horizon scientific workflows, including spatial biology claim recovery, biomedical machine-learning pipeline construction, biosecurity-relevant biological capabilities, and ontology curation for natural phenotypes. ^23,24,28,29^ However, the types of open-ended, multi-stage scientific analyses that are common to real-world research and industrial applications remain underexamined.

GeneBench-Pro updates our previously released GeneBench eval with substantially difficulty-hardened versions of the original problems and new problems in new domains. It uses a tiered release model rather than a single fully public 129-problem repository. Ten complete, externally reviewed problem packages are publicly available on Hugging Face, including solver-facing prompts, staged data, graders, and detailed reports. Fifty additional held-out problems were provided to Artificial Analysis for independent third-party model benchmarking, with priority given to relatively difficult tasks that remain informative for third-party evaluation. The remaining 69 problems are retained as an internal holdout to support continued evaluation while reducing the risk of benchmark contamination.

In our evaluations, the strongest models continue to show substantial partial competence across many tasks, even when they do not complete the full decision-making chain. We observe that while frontier models consistently notice data issues, statistical irregularities, and other potential problems, there remains an incomplete ability to bridge the “notice-act” gap required to close the inferential loop. Qualitatively, this pattern resembles expert-novice differences in scientific problem solving observed in humans, where experts utilize their experience to guide problem representation and adaptive decision-making, while novices struggle to integrate observations into the broader context of the problem. ^30,31^ We therefore anticipate that improvements in planning, self-revision, and uncertainty-aware control should translate into meaningful gains on this class of work. ^32–34^

Realizing these capability gains depends on having evaluations that can reliably measure progress; while GeneBench-Pro improves upon GeneBench in evaluating this gap in capabilities, it shares similar limitations. Constructive staging and simulation make the endpoint identifiable and the grading interpretable, but GeneBench-Pro does not attempt to reproduce the documentation gaps, data scale, and study-specific irregularities of true real-world analyses. ^35^

The all-or-nothing nature of our binary grading is an explicit measurement choice to reflect analogous real-world situations; in a scientific or translational workflow, an agent that executes several intermediate steps correctly but returns the wrong decision-relevant answer has not successfully automated the analysis. At the same time, this collapses useful stage-level diagnostic evidence, since a run that resolves most decision points but fails late is scored the same as one that fails immediately. Future versions of GeneBench-Pro may therefore add auxiliary stage-level or rubric-based scoring to measure partial progress, while retaining end-to-end pass rate as the primary metric.

Enabling agents to reliably automate this class of analysis could significantly accelerate scientific discovery. For example, human genetic evidence has played an increasingly central role in target prioritization and translational follow-up, ^36^ where mechanisms with human genetic support are materially more likely to translate into approved indications. ^37–39^ The plummeting costs of sequencing and the expansion of biobank-scale resources with linked molecular, phenotype, and health record data have enabled this trend to accelerate, ^40–43^ but one of its consequences is that the bottleneck is increasingly shifting from data generation to the ability to turn data into actionable insights.

Models that could consistently execute the types of analyses that currently require teams of expert analysts would therefore have a transformative impact on the throughput and nature of industrial research by accelerating hypothesis triage, target follow-up, and the iteration cycle between data generation and decision-making. As a rough point of reference, executed unaided by a human expert, a typical GeneBench-Pro problem would take on the order of 10–40 hours all-in. At a conservative $100–$200 per hour, the human labor cost of a single problem is already on the order of a few thousand dollars. Current frontier-agent attempts are still too unreliable to replace that labor, but the cost asymmetry implies that even partial automation could be operationally valuable if models become able to close the remaining inferential loop. These figures are only illustrative, but they indicate that the value of reliable automation on tasks of this type could be substantial even before considering the effects of scale or accelerated iteration speed.

Our results indicate that while current models continue to make substantial progress toward automating these analyses, there remains a significant capabilities gap that separates current frontier models from the reliable end-to-end performance required to fulfill this potential.

## Methods

### Evaluation and grading

Evaluation was conducted on the full 129-problem GeneBench-Pro suite. The result table includes 60 evaluated model configurations spanning GPT-5.2, GPT-5.4, GPT-5.5, GPT-5.6 Luna/Terra/Sol, GPT Pro (Extended) runs, Claude Opus, Gemini, Grok, GLM, Kimi, DeepSeek, MiMo, Tencent, MiniMax, and Qwen models. Rows are listed in **Supplementary Table 1**. For each model–problem pair, we ran 10 independent attempts for standard evaluations and 5 independent attempts for GPT Pro (Extended) and Claude Opus evaluations; attempts ending in container, tooling, provider, or response-format errors were excluded from per-problem pass-rate estimates. The final set of valid attempts for each model–problem pair was used to compute per-problem pass rates. Confidence intervals for pass rates were computed by hierarchical bootstrap, resampling problems and repeated runs within each sampled problem.

Average-token values in **Figure 4C** are reported only for mainline GPT-family rows, for which comparable trace and response-token accounting was available.

Runs used the same harness as that previously described in GeneBench: a Linux environment in a Docker container with Python and R scientific computing libraries. Relevant installed tooling included standard numerical, statistical, plotting, and file-I/O packages such as `numpy`, `pandas`, `scipy`, `scikit-learn`, `statsmodels`, `lifelines`, `matplotlib`, `seaborn`, and `openpyxl`; genomics and bioinformatics tools and libraries including PLINK 2.0, `bedtools`, `tabix`, `pysam`, `cyvcf2`, `pybedtools`, `pybigwig`, `pyfaidx`, `pysnptools`, `bed-reader`, and `pyranges`; and single-cell and expression-analysis tooling including `scanpy`, `anndata`, `scrublet`, `PyDESeq2`, `Seurat`, `DESeq2`, `edgeR`, `limma`, `tximport`, `SingleCellExperiment`, `zellkonverter`, and `scDblFinder`. The execution environment had no internet access; agents were limited to the prompt, staged files, installed software, and model-internal knowledge.

For each problem, the model is supplied with a series of initial instructions in the following order:

- a brief system message describing the container execution environment,
- the content of the prompt specifying the question at hand,
- instructions to return the final answer in a prespecified JSON schema including both numerical estimates and a brief, free-form summarization of its reasoning, and
- an enumeration of the locally mounted locations of all relevant data files.

Binary grading was performed based on pre-specified problem-specific target fields, exact-match rules, and absolute numeric tolerances. A run is counted as passing only if all graded fields satisfied their respective constraints. We report pass rates over repeated runs as the primary benchmark metric and do not use partial-credit or diagnostic scoring pathways for the primary metric. A free-text reasoning field is also collected for qualitative analysis but is not graded. Model responses were automatically graded by Python scripts encoding these constraints.

A small minority of runs (fewer than 1%) with invalid execution traces due to container, tooling, provider, or response-format failures were excluded from analysis. Models were not subject to an additional uniform wall-clock budget imposed by our harness; runs remained subject to provider and platform behavior in the evaluation stack.

## Data availability

The public GeneBench-Pro release is available through a Hugging Face dataset record. The public release contains ten open-source problems, including solver-facing prompts, staged data files, grading/configuration files, and report PDFs. These report PDFs describe the problem setup, construction, validation evidence, and grading contract, and serve as the detailed case studies for the public release. They are available within the corresponding public problem packages. The 50 Artificial Analysis problems and the remaining internal holdout are not publicly released.

## Statement on AI use

AI systems were used to assist with the design and review of problems and editing of the manuscript. The authors reviewed and approved the benchmark design, problem specifications, validation targets, evaluation results, and final manuscript text.

## Author contributions

J.L. conceived of GeneBench-Pro, implemented and reviewed problems, coordinated the external reviewers, and performed benchmark testing. A.H. ran the model evaluations. All authors contributed to drafting and revising the manuscript.

## Acknowledgements

We thank Dylan Steinecke, Toby Baker, Joseph Pickrell and Joy Jiao for helpful discussions and feedback on earlier drafts of this manuscript. We also thank the external reviewers who submitted external reviews of GeneBench-Pro problems: Toby Baker, Muhammad Elsadany, Favour N. Esedebe, Lex Flagel, David Gibson, Jennifer Grundman, Tuomo Kiiskinen, Dylan Steinecke, Zhixin Cyrillus Tan, Alex Strudwick Young, and Nicole Zeltser.

---

**Supplementary Figure 1:**
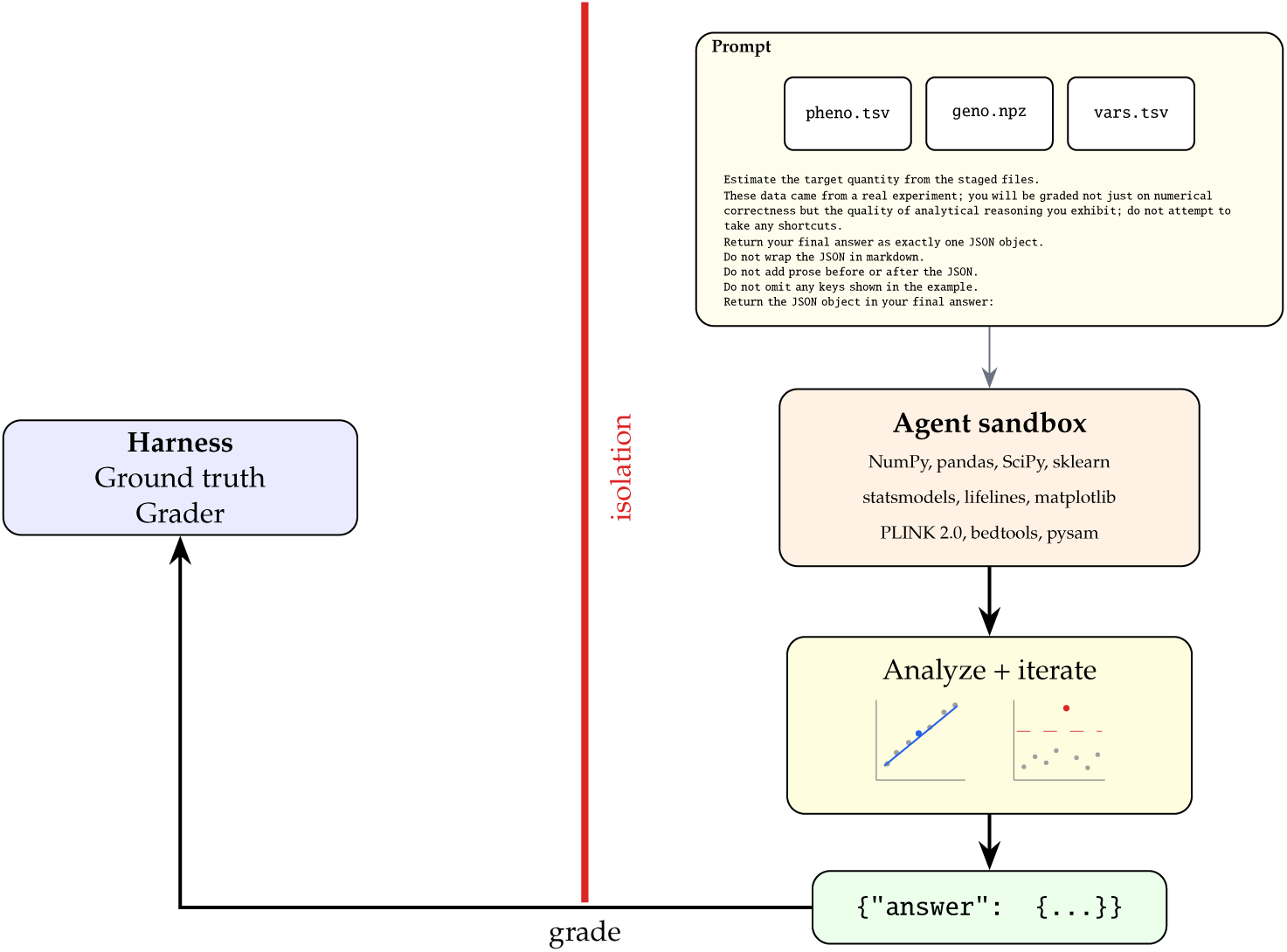
Agent environment and GeneBench-Pro problem anatomy. An agent receives a prompt, a set of files in an isolated workspace, general-purpose scientific libraries, and standard genomics bioinformatics tools. It must explore the data, test hypotheses, and execute an analysis before producing a final estimate of the target quantity in JSON format for grading.

**Supplementary Table 1:**
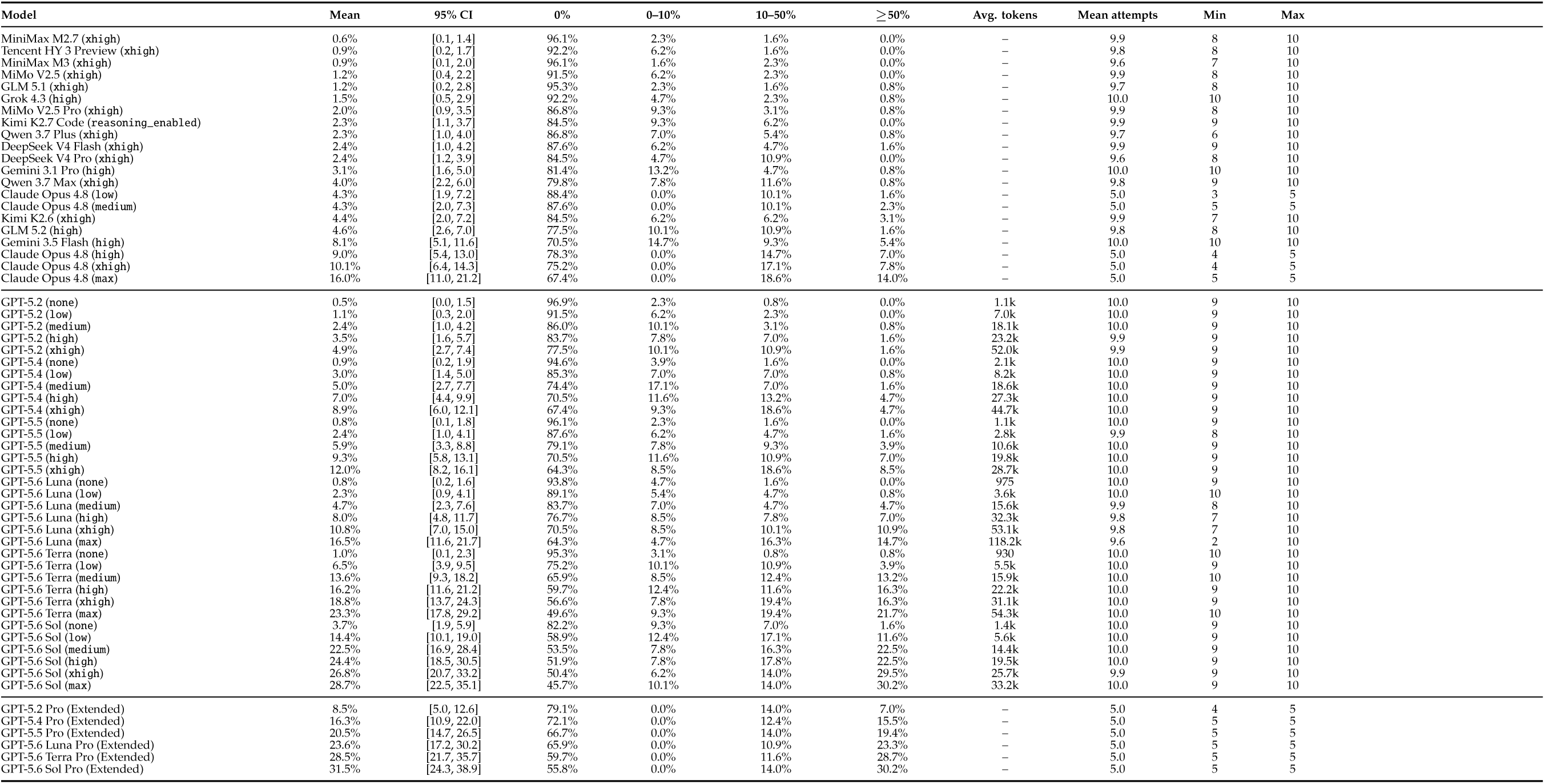
Values underlying Figure 4. Overall pass rate is the unweighted mean of per-problem pass rates across the 129 GeneBench-Pro problems. The 95% confidence intervals match **Figure 4A** and are hierarchical bootstrap intervals that resample problems and repeated runs within each problem. The regime columns match **Figure 4B**. Average-token values are shown for mainline GPT-family rows, where comparable trace and response-token accounting was available. Attempt summaries report the mean, minimum, and maximum numbers of valid attempts contributing to each model–problem pass rate after filtering error samples.

**Supplementary Table 2:**
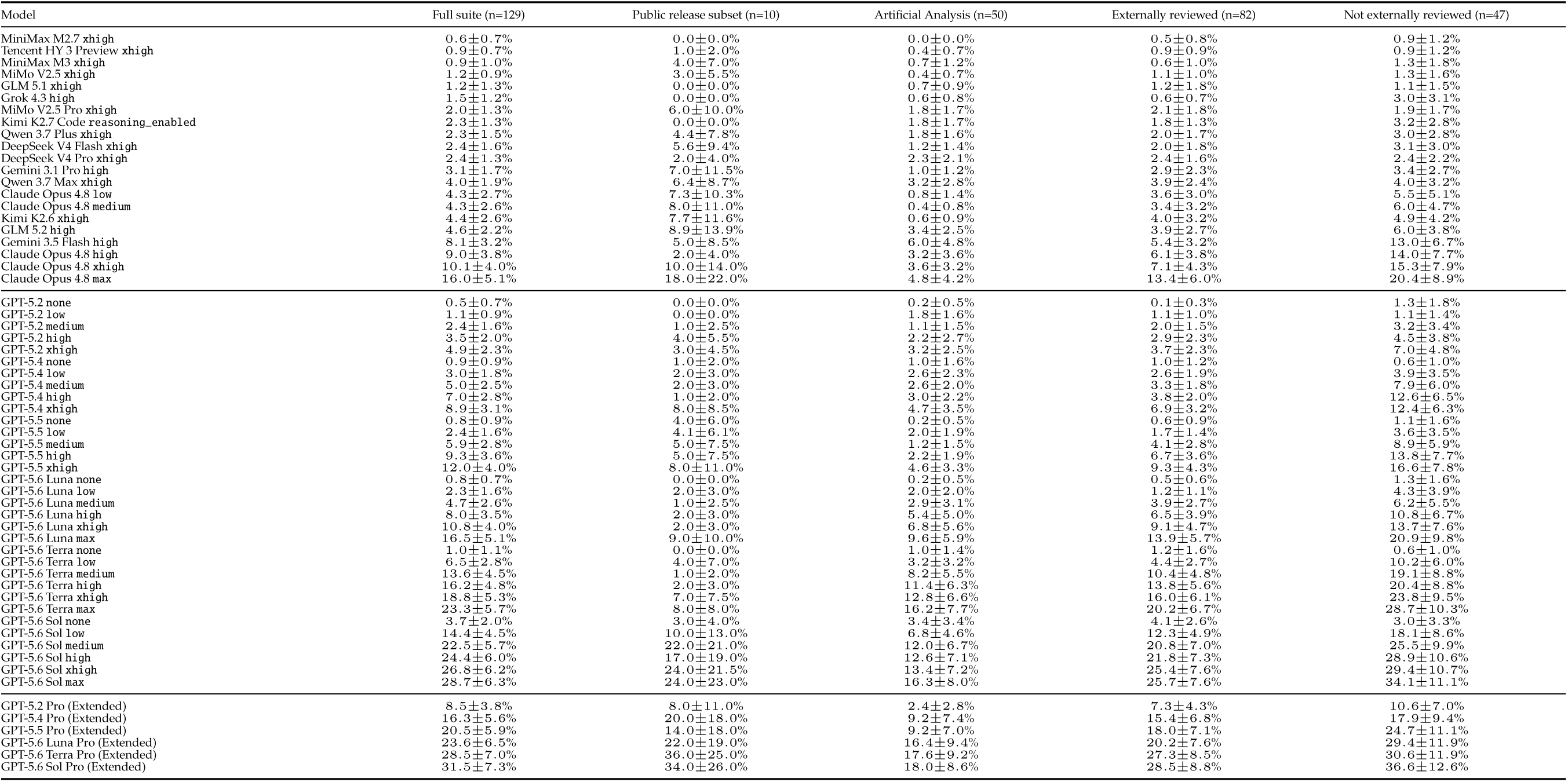
Pass rates by release subset and external review status. Rows include every evaluated model configuration. Values are unweighted means of per-problem pass rates after aggregating repeated runs within each problem; uncertainty is reported as the half-width of a 95% hierarchical bootstrap confidence interval from 20,000 resamples, resampling problems and repeated runs within each problem. The public release subset (n=10) reflects the final public case-study set; the Artificial Analysis subset (n=50) follows the final AA reporting set. The review-status columns are restricted to the 129-problem evaluation suite (82 externally reviewed; 47 not externally reviewed).

